## supplemental figures for "Mutations that prevent phosphorylation of the BMP4 prodomain impair proteolytic maturation of homodimers leading to lethality in mice"

Supplementary information included:

Figures S1-S7

Table S1-S2

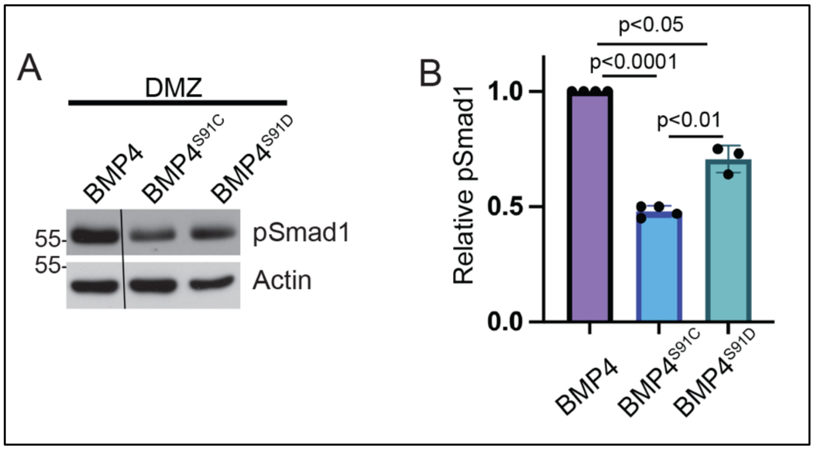


**Figure S1. Ectopic expression of a phosphomimetic mutant form of BMP4 (BMP4$^{S91D}$) generates more activity than BMP4$^{S91C}$ but less activity than native BMP4.** (A) RNA encoding a wild type or point mutant form of BMP4 (25 pg) was injected near the dorsal marginal zone (DMZ) of 4-cell embryos. DMZ explants were isolated at stage 10 and pSmad1 levels analyzed by immunoblot. Blots were reprobed with actin as a loading control. Black bar indicates where a non-relevant intervening lane was removed using photoshop. (B) Quantitation of relative pSmad1 levels normalized to actin in at least 3 independent experiments (mean +/- SD, data analyzed using an unpaired t-test).

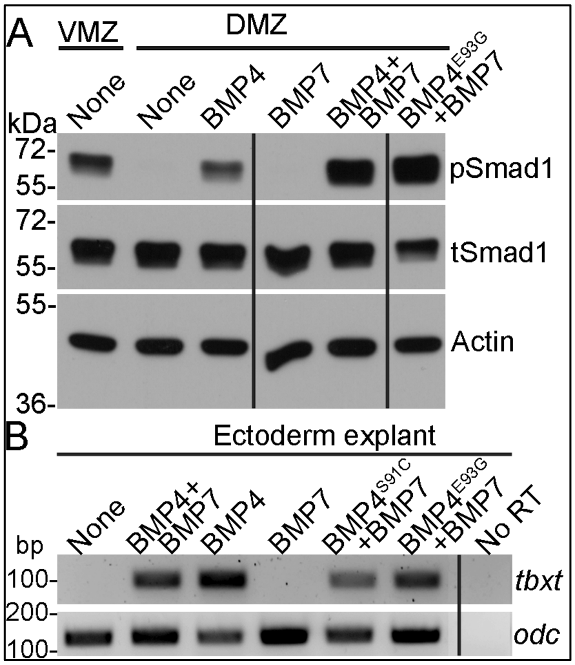


**Figure S2. Point mutations predicted to interfere with phosphorylation of the BMP4 prodomain do not interfere with BMP4/7 heterodimer activity.** (A) RNA encoding wild type BMP4 or BMP7 alone (25 pg), or one half dose wild type or point mutant BMP4 together with BMP7 (12.5 pg each) were injected near the dorsal marginal zone (DMZ) of 4-cell embryos. DMZ and ventral marginal zone (VMZ) explants were isolated at stage 10 and pSmad1 levels were analyzed by immunoblot. Duplicate blots were probed with actin as a loading control. (B) RNA encoding wild type BMP4 or BMP7 alone (25 pg), or one half dose wild type or point mutant forms of BMP4 together with BMP7 (12.5 pg each) were injected near the animal pole of two-cell embryos. Ectodermal explants were isolated at stage 10 and *tbxt* levels were analyzed by semi-quantitative RT-PCR. Black bars indicate where a non-relevant intervening lane was removed using photoshop.

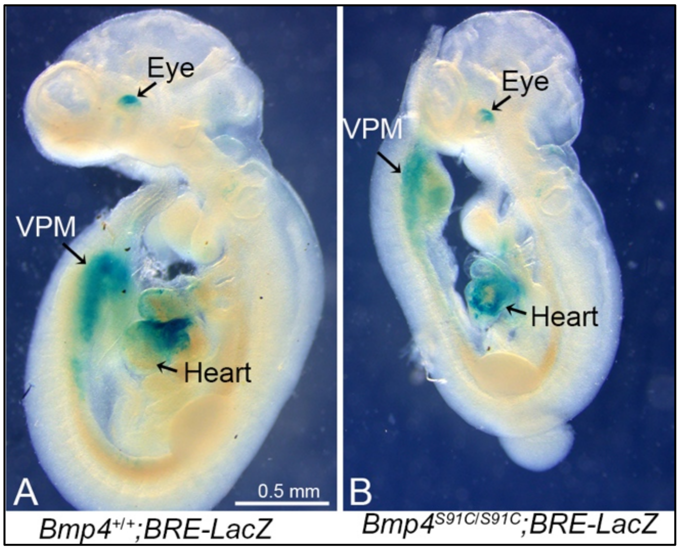


**Figure S3. Bmp4$^{S91C}$ homozygotes show reduced BMP activity in the heart and ventral posterior mesoderm at E9.5.** (A, B) E9.5 wild type (A) or *Bmp4$^{S91C/S91C}$* mutant littermates (B) carrying a BRE-LacZ transgene were stained for ß-galactosidase activity to detect endogenous BMP pathway activation. Embryos from the same litter were stained for an identical time under identical conditions. Three embryos of each genotype were examined and results shown were reproduced. VPM; ventral posterior mesoderm.

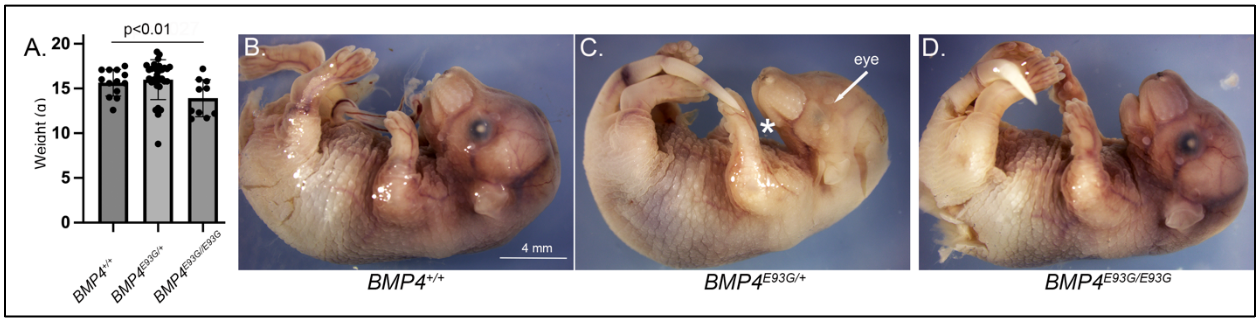


**Figure S4. *Bmp4$^{E93G}$* homozygotes are smaller than wild type littermates and a subset of mutants have eye and/or craniofacial defects.** (A) Weights of male wild type and *Bmp4$^{E93G}$* heterozygous and homozygous mutant littermates at P28 (mean +/- SD, data analyzed using an unpaired t-test). (B-D) Photographs of E17.5 wild type and mutant littermates. A subset of *Bmp4$^{E93G}$* heterozygotes show craniofacial defects such as a small mandible (asterisk) and small or absent eyes (arrow) at low frequency (*n*=1/12).

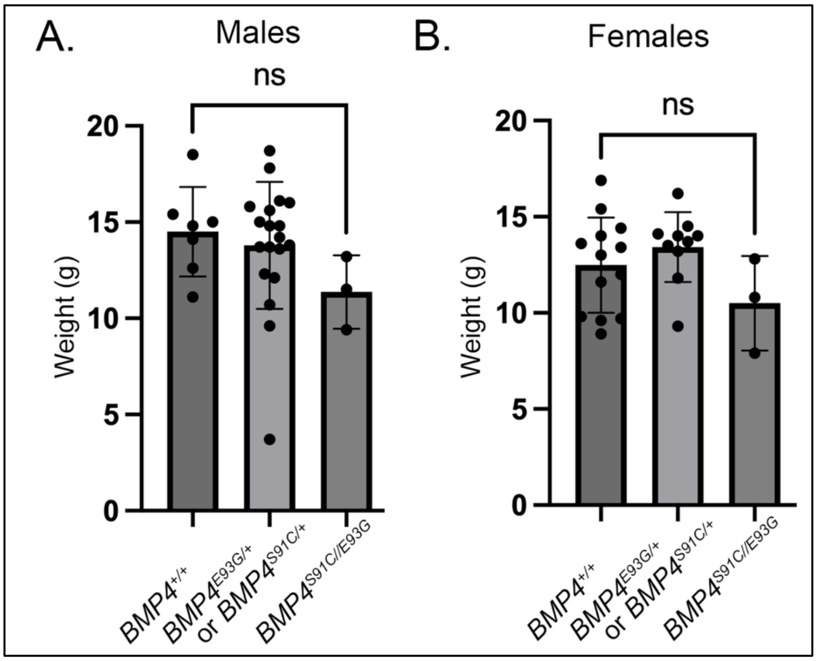


**Figure S5. *Bmp4*$^{S91C/E93G}$ compound heterozygotes are slightly smaller than heterozygous or wild type littermates.** (A, B) Weights of male (A) and female (B) wild type, heterozygous and compound heterozygous mutant littermates at P28 (mean +/- SD, data analyzed using an unpaired t-test). Genotyping protocol does not distinguish between the *Bmp4*$^{S91C/+}$ or *Bmp4*$^{E9G3G/+}$ allele and thus weights of *Bmp4*$^{S91C/+}$ or *Bmp4*$^{E9G3G/+}$ heterozygotes are reported together.

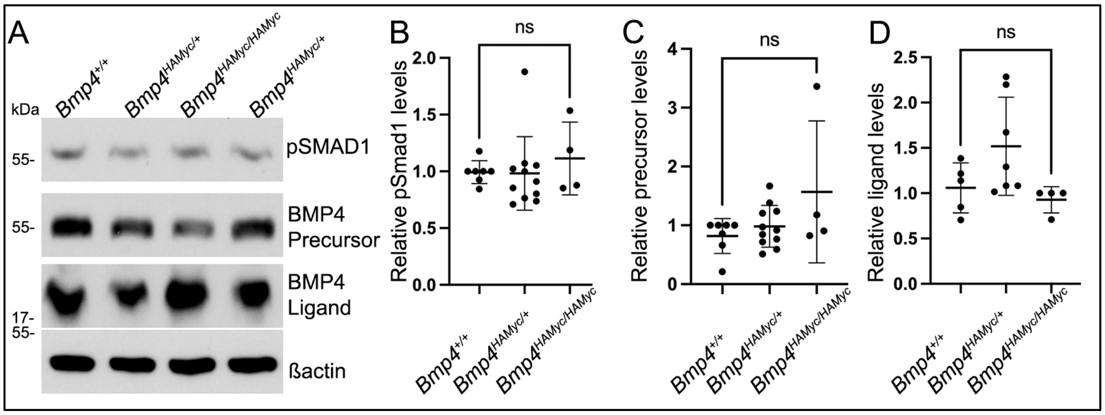


**Figure S6. Levels of BMP4 precursor protein, cleaved ligand and pSMAD1 are unchanged in MEFs isolated from Bmp4$^{+/+}$ embryos carrying epitope tags relative to untagged littermates.** (A-D) Levels of pSMAD1, BMP4 precursor protein or cleaved BMP4 ligand were analyzed in MEFs isolated from *Bmp4$^{HAMyc}$* embryos or wild type littermates at E13.5. A representative blot (A) and quantitation of protein levels normalized to actin (mean +/- SD) (B-D) are shown.

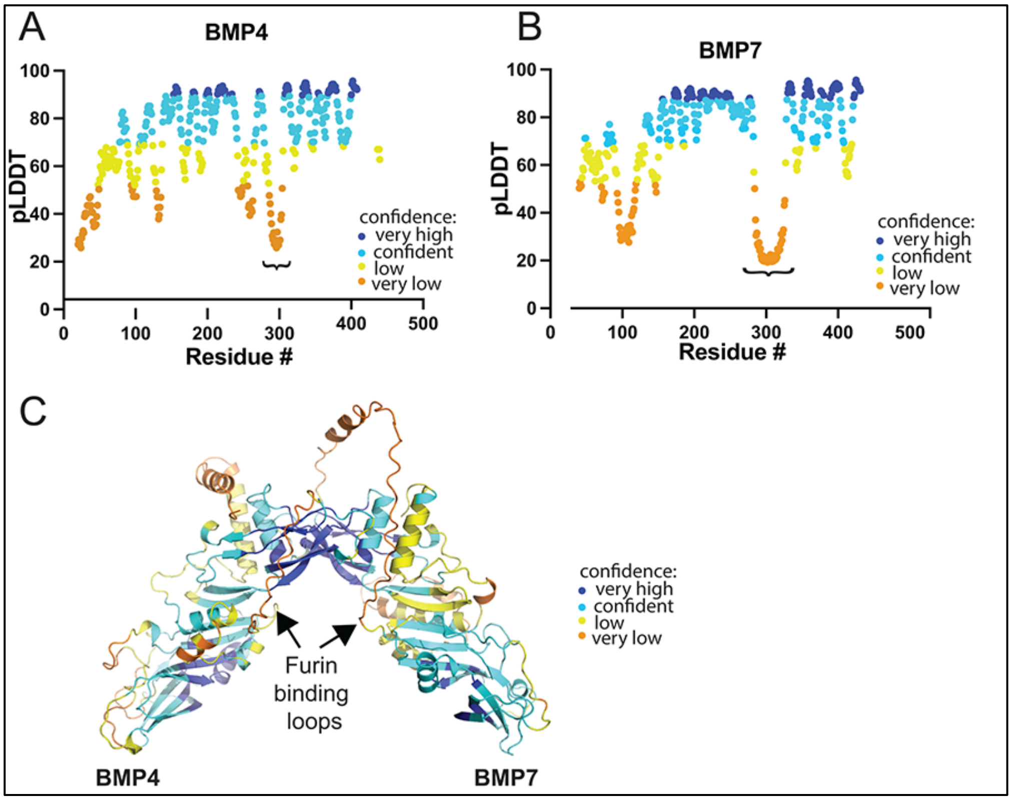


**Figure S7. Relative confidence in structural predictions across different regions of BMP4 and BMP7 precursor proteins.** (A-C) Bmp4 and Bmp7 precursor proteins were modeled in Alphafoldserver.com (version 3). Representative predictions are shown colored according to relative confidence (pLDDT) at the level of amino acid residues from N- to C-terminus in BMP4 (A) and BMP7 (B) monomers and in the predicted structure of BMP4/7 heterodimers (C). The furin binding motifs (denoted by "} "in A, B; arrows in C) lie within a poorly predicted region, but the location of that region with respect to the overall structure is reproducible across multiple types of predictions (homodimer, heterodimer, pSer-modified dimers).

**Table S1. Progeny from *Bmp4*$^{S91C/+}$ intercrosses at embryonic stages by sex.**

**A. Progeny from *Bmp4*$^{S91C/+}$ intercrosses at E13.5**

| Sex | *Bmp4*$^{+/+}$ | *Bmp4*$^{S91C/+}$ | *Bmp4*$^{S91C/S91C}$ | *n* | *p* |
|---|---|---|---|---|---|
| Both | 22 (19) | 55 (38) | 0 (19) | 77 | 0.00001 |
| Male | 12 (11) | 32 (22) | 0 (11) | 44 | 0.00015 |
| Female | 10 (8) | 23 (17) | 0 (8) | 33 | 0.00131 |

**B. Progeny from *Bmp4*$^{S91C/+}$ intercrosses at E11.5**

| Sex | *Bmp4*$^{+/+}$ | *Bmp4*$^{S91C/+}$ | *Bmp4*$^{S91C/S91C}$ | *n* | *p* |
|---|---|---|---|---|---|
| Both | 23 (16) | 39 (31) | 0 (16) | 62 | 0.00001 |
| Male | 15 (9) | 22 (19) | 0 (9) | 37 | 0.00118 |
| Female | 8 (6) | 17 (13) | 0 (6) | 25 | 0.01530 |

**C. Progeny from *Bmp4*$^{S91C/+}$ intercrosses at E10.5**

| Sex | *Bmp4*$^{+/+}$ | *Bmp4*$^{S91C/+}$ | *Bmp4*$^{S91C/S91C}$ | *n* | *p* |
|---|---|---|---|---|---|
| Both | 58 (60) | 135 (121) | 46 (60) | 239 | 0.072 |
| Male | 34 (33) | 69 (67) | 28 (33) | 131 | 0.630 |
| Female | 24 (27) | 66 (54) | 18 (27) | 108 | 0.065 |

**D. Progeny from *Bmp4*$^{S91C/+}$ intercrosses at E9.5**

| Sex | *Bmp4*$^{+/+}$ | *Bmp4*$^{S91C/+}$ | *Bmp4*$^{S91C/S91C}$ | *n* | *p* |
|---|---|---|---|---|---|
| Both | 7 (12) | 31 (24) | 9 (12) | 47 | 0.084 |
| Male | 4 (7) | 21 (15) | 4 (7) | 29 | 0.054 |
| Female | 3 (5) | 10 (9) | 5 (9) | 18 | 0.72 |

(A-D) Numbers of observed and expected (in parenthesis) embryos of each genotype listed in the top row are indicated. The p value is based on X2 test.

**Table S2. Progeny from *Bmp4*$^{E93G/+}$ x *Bmp4*$^{-/+}$ crosses at embryonic stages by sex.**

**A. Progeny from *Bmp4*$^{E93G/+}$ x *Bmp4*$^{-/+}$ crosses by sex at E13.5-14.5**

| Sex | *Bmp4*$^{+/+}$ | *Bmp4*$^{E93G/+}$ | *Bmp4*$^{-/+}$ | *Bmp4*$^{-/E93G}$ | *n* | *p* |
|---|---|---|---|---|---|---|
| Both | 17 (17) | 13 (17) | 19 (17) | 17 (17) | 66 | 0.765 |
| Male | 10 (9) | 6 (9) | 9 (9) | 10 (9) | 35 | 0.746 |
| Female | 7 (8) | 7 (8) | 10 (8) | 7 (8) | 31 | 0.832 |

**B. Progeny from *Bmp4*$^{E93G/+}$ x *Bmp4*$^{-/+}$ crosses by sex at E11.5-12.5**

| Sex | *Bmp4*$^{+/+}$ | *Bmp4*$^{E93G/+}$ | *Bmp4*$^{-/+}$ | *Bmp4*$^{-/E93G}$ | *n* | *p* |
|---|---|---|---|---|---|---|
| Both | 11 (10) | 4 (10) | 12 (10) | 14 (10) | 41 | 0.624 |
| Male | 6 (6) | 2 (6) | 8 (6) | 7 (6) | 23 | 0.307 |
| Female | 5 (5) | 2 (5) | 4 (5) | 7 (5) | 18 | 0.409 |

**C. Progeny from *Bmp4*$^{E93G/+}$ x *Bmp4*$^{-/+}$ crosses by sex at E10.5**

| Sex | *Bmp4*$^{+/+}$ | *Bmp4*$^{E93G/+}$ | *Bmp4*$^{-/+}$ | *Bmp4*$^{-/E93G}$ | *n* | *p* |
|---|---|---|---|---|---|---|
| Both | 7 (7) | 10 (7) | 7 (7) | 5 (7) | 29 | 0.136 |
| Male | 3 (3) | 4 (3) | 4 (3) | 2 (3) | 13 | 0.838 |
| Female | 4 (4) | 6 (4) | 3 (4) | 3 (4) | 16 | 0.682 |

(A-C) Numbers of observed and expected (in parenthesis) embryos of each genotype listed in the top row are indicated. The p value is based on X2 test.
